# Palatal segment contributions to midfacial anterior-posterior growth

**DOI:** 10.1101/2023.10.03.560703

**Authors:** Ian C. Welsh, Maria E. Feiler, Danika Lipman, Isabel Mormile, Karissa Hansen, Christopher J. Percival

## Abstract

Anterior-posterior (A-P) elongation of the palate is a critical aspect of integrated midfacial morphogenesis. Reciprocal epithelial-mesenchymal interactions drive secondary palate elongation that is coupled to the periodic formation of signaling centers within the rugae growth zone (RGZ). However, the relationship between RGZ-driven morphogenetic processes, the differentiative dynamics of underlying palatal bone mesenchymal precursors, and the segmental organization of the upper jaw has remained enigmatic. A detailed ontogenetic study of these relationships is important because palatal segment growth is a critical aspect of normal midfacial growth, can produce dysmorphology when altered, and is a likely basis for evolutionary differences in upper jaw morphology. We completed a combined whole mount gene expression and morphometric analysis of normal murine palatal segment growth dynamics and resulting upper jaw morphology. Our results demonstrated that the first formed palatal ruga (ruga 1), found just posterior to the RGZ, maintained an association with important nasal, neurovascular and palatal structures throughout early midfacial development. This suggested that these features are positioned at a proximal source of embryonic midfacial directional growth. Our detailed characterization of midfacial morphogenesis revealed a one-to-one relationship between palatal segments and upper jaw bones during the earliest stages of palatal elongation. Growth of the maxillary anlage within the anterior secondary palate is uniquely coupled to RGZ-driven morphogenesis. This may help drive the unequaled proportional elongation of the anterior secondary palate segment prior to palatal shelf fusion. Our results also demonstrated that the future maxillary-palatine suture, approximated by the position of ruga 1 and consistently associated with the palatine anlage, formed predominantly via the posterior differentiation of the maxilla within the expanding anterior secondary palate. Our ontogenetic analysis provides a novel and detailed picture of the earliest spatiotemporal dynamics of intramembranous midfacial skeletal specification and differentiation within the context of the surrounding palatal segment A-P elongation and associated rugae formation.

## Introduction

Morphological variation of the midfacial complex, which consists of the nose, upper jaw, cheek, and palate, is a defining aspect of both intra- and inter-specific differences in facial shape. Facial shape morphogenesis begins with the specification of cranial neural crest (CNC) cells during neurulation, which then delaminate around mouse embryonic day (E) 8.5 and migrate to drive facial prominence outgrowth between E9 and E10 (Minoux and Rijli, 2010; Dash and Trainor, 2020). By mouse E11.5, fusion between the medial nasal processes (mnp) of the frontonasal process (FNP) and the anterior maxillary processes (MxP) gives rise to the primary palate and lip, producing a unified upper jaw from previously separated tissues (Fig. 1).

**Figure 1.**
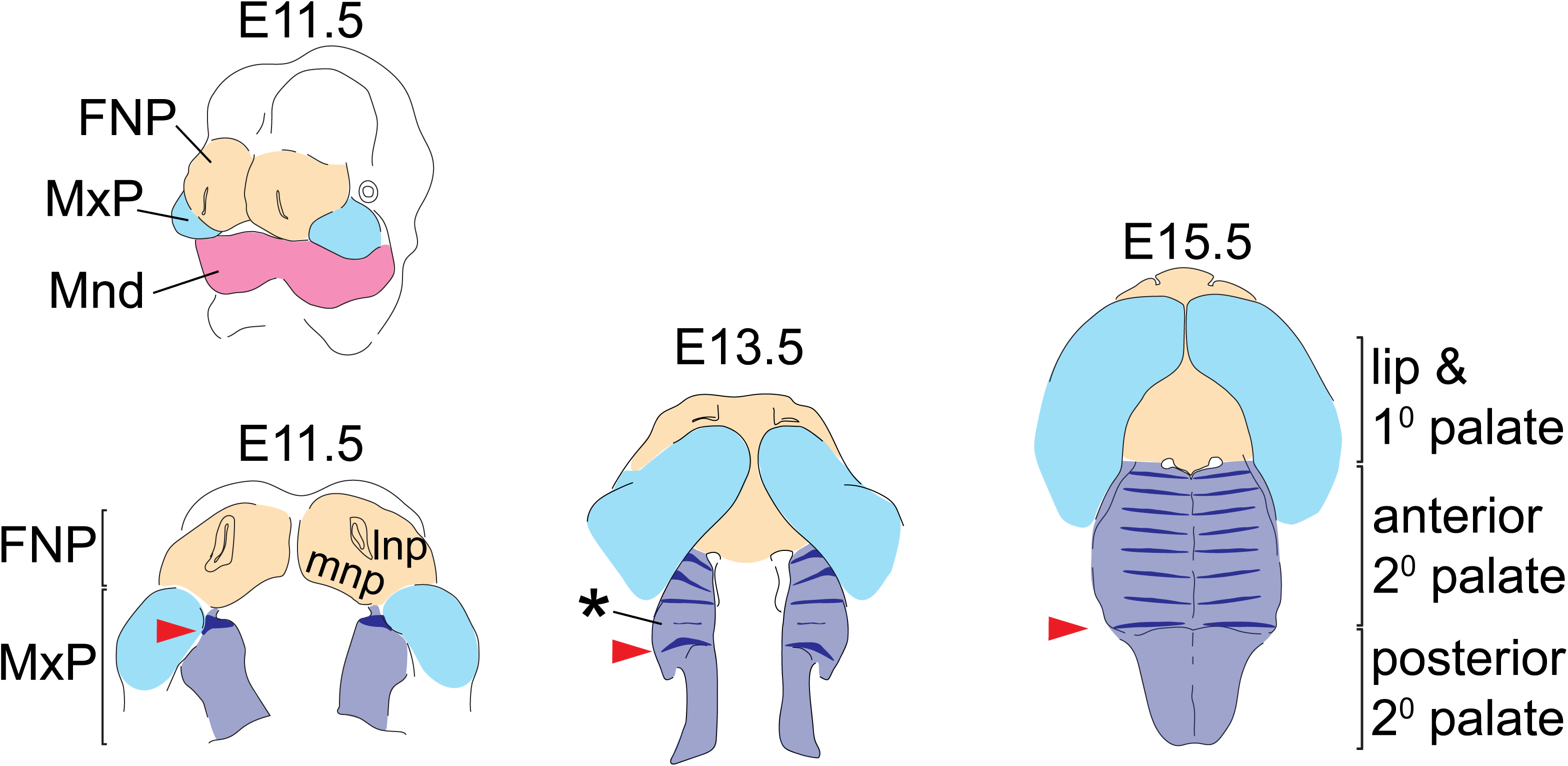
Tissue origins of the midfacial complex and rugae position during secondary palate morphogenesis – The upper and lower jaws are formed from the frontonasal process (FNP - tan), and branchial arch 1 derived maxillary and mandibular processes (MxP - pale blue and Mnd – magenta, respectively). From E11.5 to E15.5, outgrowth and fusion of the medial nasal process (mnp) with the superficial portion of the MxP frame out the lip and primary (1°) palate, while the A-P elongation and medially directed growth of the palatal shelves from the internal portion of the MxP gives rise to the secondary palate (2° pal – light purple). *Shh* expression (dark blue) highlights the dynamics of rugae formation and illustrates regional expansion of the anterior secondary palate. At E11.5, ruga 1 (red arrowheads) forms at the anterior extent of the nascent palatal shelf and subsequently defines the caudal end of the rugae growth zone (asterisk) where new rugae form prior to being displaced anteriorly. Additional abbreviation: lateral nasal process (lnp).

Secondary palate morphogenesis then begins through outgrowth of the nascent palatal shelf along the medial aspect of the MxP. Significant palatal growth along the anterior-posterior (A-P) axis is accompanied by vertical growth of the palatal shelves prior to their elevation and medial fusion dorsal to the tongue, after which the palate separates the oral and nasal cavities (Bush and Jiang, 2012; Hammond and Dixon, 2022). Disproportional growth of the FNP or MxP prior to fusion can result in cleft lip and palate (Young et al., 2014), while poor coordination between the multiple growth axes of MxP-derived tissues can prevent secondary palate closure even when the palatal shelves maintain competency to fuse (Kouskoura, et al., 2013).

The three longitudinal segments of the palate are called the primary palate, anterior secondary palate, and posterior secondary palate (Fig. 1). The boundary between primary palate and anterior secondary palate is the posterior edge of the fused mnp. Although A-P organization of the secondary palate can be defined by the presence of boney versus muscular tissue (i.e., the hard versus soft palate) or modes of palatal shelf elevation and closure (reviewed by Yu and Ornitz, 2011), we define secondary palate segments by molecular differences in tissue patterning and cell signaling competence (Hilliard et al., 2005; Hammond and Dixon, 2022).

The A-P expression domains of certain transcription factors (TFs) and intercellular signaling molecules within CNC-derived mesenchyme are important for A-P organization and growth during palatal morphogenesis. For example, within this mesenchyme, *Msx1* and *Shox2* are exclusively expressed in the anterior secondary palate while *Barx1* and *Tbx22* are exclusive to the posterior secondary palate (Hilliard et al., 2005; Okano et al., 2006; Welsh et al., 2018). Mutations of *Shox2* or *Tbx22* have resulted in midline clefts of the anterior or posterior palate, respectively (Yu et al., 2005; Pauws et al., 2009), although regional expression is maintained in genetically engineered mice with severe segmental growth defects (Welsh et al., 2018). The A-P expression boundary of these and other factors is found at the first formed palatal ruga (Hilliard et al., 2005; Li and Ding, 2007; Welsh et al., 2007; Welsh and O’Brien, 2009; Welsh et al., 2018; Hammond and Dixon, 2022), one of multiple rugae, which are parallel epithelial thickenings on the anterior secondary palate.

This first ruga (numbered according to Welsh et al., 2007, Welsh and O’Brien, 2009; although Pantalacci et al., 2008 use a different numbering system) forms at E11.5 on the anterior extent of the nascent secondary palate (red arrow in Fig. 1). Remaining rugae form sequentially within the rugae growth zone (RGZ) located just anterior to ruga 1 (Welsh and O’Brien, 2009), via a Turing type mechanism (Economou et al., 2012; Kawasaki et al., 2018). Rugae are centers of Sonic Hedgehog (SHH) signaling, which is critical for anterior secondary palate patterning and A-P growth. As the secondary palate completes midline fusion at E15.5, the last formed ruga appears adjacent to ruga 1 in the middle of the secondary palate while each precedingly formed ruga is found at an increasingly anterior position along the anterior secondary palate (Welsh and O’Brien, 2009).

Based upon the dynamics of rugae formation, palate growth, and patterning gene expression, Pantalacci (et al., 2008) hypothesized that ruga 1 represents a distinct morphological boundary related to either the future hard versus soft palate boundary or the maxillary-palatine suture. We formally test these alternate hypotheses within our analysis of tissue relationships and cell identity markers across a developmental time course spanning mouse secondary palate morphogenesis and upper jaw outgrowth (E11.5-E15.5). More broadly, our analysis explores the degree to which rugae formation and regional A-P patterns of secondary palate gene expression are coordinated with palate skeletal differentiation and elongation across this critical developmental period.

Our results establish one-to-one palatal segment origins of upper jaw bones, highlighting a previously unappreciated coupling of maxillary bone formation to rugae morphogenesis during initial elongation of the anterior secondary palate (E11-E15). Strikingly, the proportional contribution of anterior secondary palate to overall palate length doubles during this ontogenetic period. Further, the one-to-one relationship between palatal segments and bones also confirms that ruga 1 is coincident with the future position of the maxillary-palatine suture, rather than the future boundary of the hard and soft palate. Overall, our multifaceted analysis of ontogenetic trends in palatal and midfacial elongation provides a novel contextual framework and developmental perspective within which to evaluate the impact of both non-pathogenic and pathogenic genetic differences on palatal segment contributions to midfacial growth and differentiation.

## Methods

### Specimen and Image Acquisition

Animal breeding, specimen collection, and tissue fixation were performed in accordance with the protocols of the University of California, San Francisco Institutional Animal Care and Use Committee under protocol approval number AN192776-01F. Mice were socially housed under a twelve-hour light-dark cycle with food and water *ad libitum*. Additional enrichment was provided when single housing was required for breeding purposes. Mice were euthanized by CO_2_ inhalation followed by cervical dislocation or decapitation. The C57BL/6J strain (RRID:IMSR_JAX:000664; Jackson Labs, Bar Harbor, ME) was chosen for analysis because it is the most widely utilized inbred strain for craniofacial genetic studies and because it was the strain originally used to train the eMOSS limb bud staging algorithm (Musy et al., 2018) that we employed as a basis for standardized developmental staging.

C57BL/6J embryos were collected between gestational days E11.5 and E15.5, as determined from copulatory plug occurrence. Ten postnatal day one (P1) specimens were collected. Embryo and P1 specimens were fixed in 4% PFA and stored in 1x PBS for micro-computed tomography (μCT) imaging or dehydrated through a graded PBST-MeOH series and stored at −20°C until use for in situ hybridization. A flowchart provides an overview of our study design (Fig. 2).

**Figure 2.**
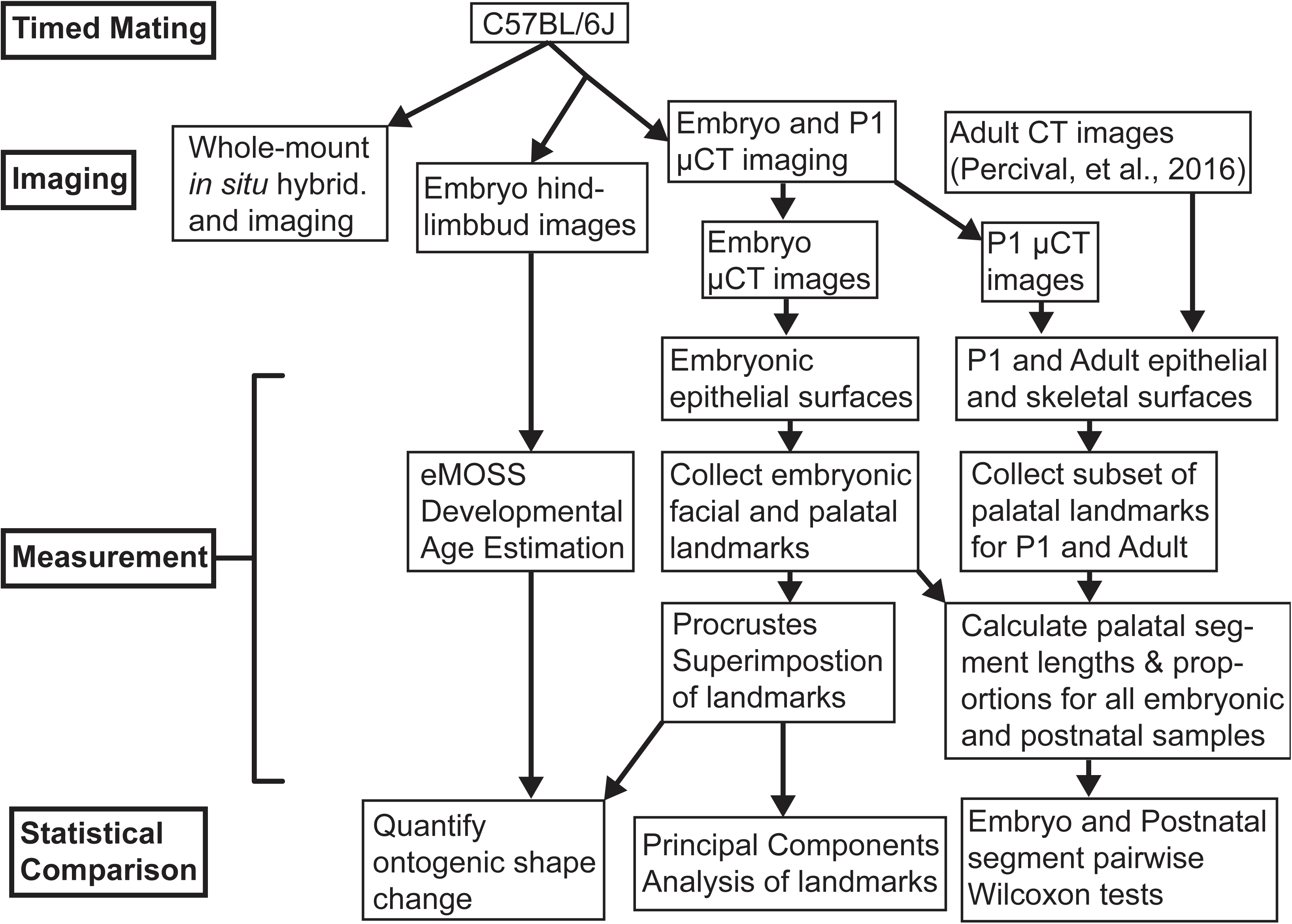
Flowchart illustrating this study’s research design.

Specimens for µCT scanning were received, stored, and imaged at the University of Calgary in accordance with the protocols of the University of Calgary Institutional Care and Use Committee under approval number AC13-0268. After approximately an hour of soaking in Cysto-Conray II (Liebel-Flarsheim Canada), embryo heads were placed upside down on cheese wax and immediately µCT scanned in air, a method that leads to minimal dehydration and good tissue surface quality of surface anatomical structures (Schmidt, et al. 2010). These µCT images were acquired with a Scanco µ35 with 45kV/177µA for images of 0.012 mm^3^ voxel size. µCT images of P1 heads were acquired similarly, but with 0.021 mm^3^ voxel size. Photographs of embryo hindlimb buds were collected using a dissecting microscope for developmental age estimation. Adult specimens were previously collected and µCT imaged as described by Percival et al., 2016.

### In Situ Hybridization

In situ hybridization was performed as described in Welsh et al., 2018. cDNA probes for *Phospho1* (nucleotides 274–1739 of RefSeq NM_153104), *Runx2* (nucleotides 1275–2704 of RefSeq NM_001271627), *Shh* (full length cDNA clone from A. McMahon, University of Southern California, Los Angeles CA, USA), *Shox2* (nucleotides 870–1603 of RefSeq NM_130458), *Sp7* (nucleotides 431–1715 of RefSeq NM_130458), and *Tbx22* (nucleotides 307– 1547 of RefSeq NM_013665) were linearized and in vitro transcribed to label with either digoxigenin (DIG) or dinitrophenol (DNP). Colorimetric detection of probes used BM purple (dark blue), BCIP (teal), or MagentaPhos (magenta). Minimally, 3-6 embryos per time point were processed for each probe analyzed.

### DMDD HREM Data

High resolution episcopic microscopy (HREM) data (Geyer et al., 2009) was generated as part of the deciphering mechanisms of developmental disorders (DMDD) program (Wilson et al., 2016) and is available from the BioImage Archive at: https://www.ebi.ac.uk/biostudies/bioimages/studies/S-BIAD490?query=DMDD.

### Embryonic Palatal Segment Growth

#### Developmental Age Estimation

Given that specimen developmental timing varies within litters and between litters of the same estimated embryonic day, we categorized embryonic specimens by developmental age rather than gestational age prior to morphometric analysis. Developmental age was estimated for each µCT scanned embryonic specimen using eMOSS, an application that predicts developmental age from hindlimb bud outlines, based on a previous analysis of C57BL/6J mice (Musy et al., 2018). The resulting limb-based estimates of developmental age were reported as days since conception, up to two decimal places. eMOSS developmental age estimates correlated strongly with the number of days after plug identification (correlation coefficient: 0.98). Limb-based developmental ages were an average of 0.26 days younger than our copulatory plug estimates. We combined similar developmental age estimates within whole- or half-day developmental age categories, which include specimens within 0.25 days of their initial eMOSS estimate (Table 1).

**Table 1.** Sample size of µCT imaged and morphometrically analyzed embryo specimens by forelimb based developmental age estimation, as estimated using eMOSS (Musy et al., 2018).

| Limb-Based Developmental Age | C57BL/6J |
| --- | --- |
| E11 | 8 |
| E11.5 | 2 |
| E12 | 7 |
| E12.5 | 4 |
| E13 | 9 |
| E13.5 | 0 |
| E14 | 26 |
| E14.5 | 1 |
| E15 | 9 |

#### Anatomical Landmark Collection

All embryo midfacial and palate landmarks were collected within Meshlab (Cignoni et al., 2008) on minimum threshold-based epithelial tissue surfaces (downsampled x2) produced from the µCT images. Landmarks that could be identified consistently in anatomically homologous positions across the embryonic period of facial growth were chosen around the nose and whisker region, the eyes, and along the palate (Fig. 3, Supporting Table 1). The midfacial landmarks we collected were previously defined to track developmentally homologous structures across multiple stages of facial development (Percival, et al., 2014) and our new set of epithelial surface palatal landmarks were defined similarly.

**Figure 3.**
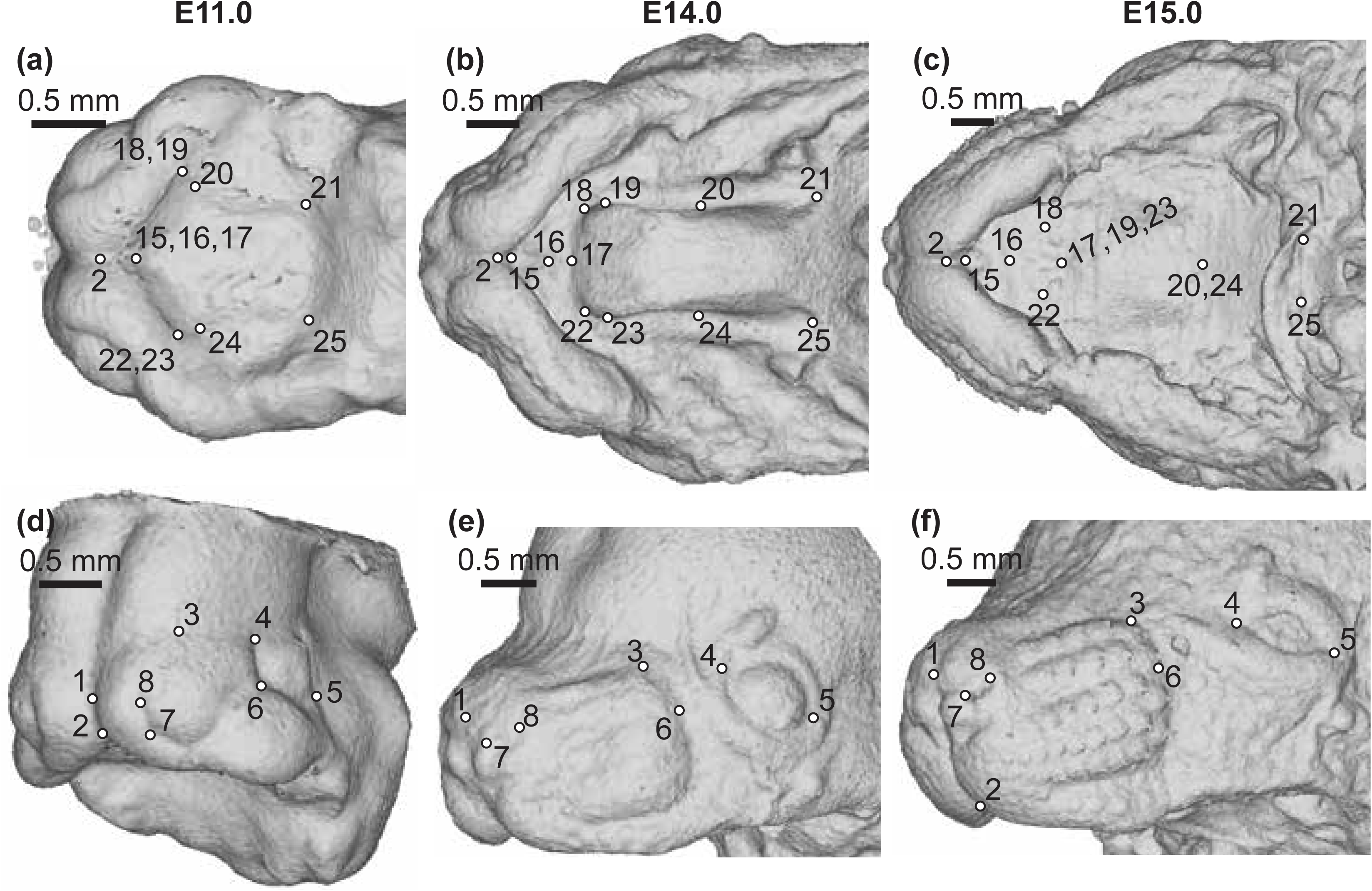
Anatomically homologous epithelial landmarks - Landmarks collected for the quantification of midfacial and palatal shape across limb based embryonic (E) developmental stages, from (a-c) palatal and (d-f) oblique facial views. Scale bars: 0.5mm. See also Supporting Table 1.

To determine if differences in proportional palate segment length contributions were consistent across embryonic and postnatal ages, a subset of epithelial palatal landmarks (Supporting Table 2) were collected on epithelial surfaces produced from ten P1 specimen CT scans. Minimum threshold-based skeletal surfaces of the same ten P1 and twenty adult (70-73 days old; 9 male and 11 female) specimens were produced using 3D Slicer (Fedorov et al., 2012) after Gaussian blur image filtering (sigma set to 0.01 for P1; sigma set to 0.02 for adult). Great care was taken to identify skeletal anatomical landmarks that closely and homologously matched the palate segment landmarks defined on surface epithelium (Supporting Table 2).

All epithelial landmarks were collected by a single observer to remove the possibility of interobserver error. All skeletal landmarks were collected by a single observer for the same reason. A reliability study was completed to quantify intraobserver error in landmark placement. Two trials were completed by the epithelial landmark observer on three E11.5, three E12.5, and three E14.5 specimens. The second trials were completed more than one year after the first trials were completed, providing a realistic estimate of error across the entire period of landmark data collection. There is no significant effect of trial or trial by age interaction on midfacial and palatal shape, indicating there is no systematic difference in how landmarks were placed between trial 1 and trial 2. The median millimeter difference in specific landmark location between trials varies across landmarks (Supporting Fig. 1), but the overall mean landmark placement difference is 0.034mm with a standard deviation of 0.022mm. This minor intraobserver error has a negligible impact on our results, given that average total palate length is 1.23mm at E11 and 2.79mm at E14.

#### Prenatal Geometric Morphometric Analysis

A Procrustes superimposition-based geometric morphometric analysis was used to quantify the ontogenetic shape change of the palate and face for embryonic specimens with limb-based developmental age estimates between E11 and E15 (Table 1). This analysis of midfacial and palatal landmark ontogeny was performed using *geomorph* (Adams et al., 2020) and RRPP (Collyer and Adams, 2018) libraries in R Statistical Software (R Core Team, 2021). Generalized Procrustes analysis (GPA) aligned specimen landmark sets by translating, scaling, and rotating their landmark coordinates (reviewed by Zelditch et al., 2012). Subsequent shape analyses were completed using the symmetric component of Procrustes-aligned specimen landmark coordinate variation, assuming that most bilateral shape differences between the left and right sides of a specimen’s face are due to random effects associated with developmental noise and tissue fixation (Palmer and Strobeck, 1986). Thus, symmetrized landmark coordinate data were interpreted to represent the midfacial and palatal shape of the C57BL/6J inbred strain genotype.

The mean midfacial shape of specimens within each developmental age category were estimated. Differences between age-specific mean shapes were plotted to illustrate typical midfacial/palatal shape ontogenetic changes during this important period of secondary palate elongation and midline fusion. A principal components analysis (PCA) was completed to identify the major axes of shape covariation across the embryonic sample.

### Palate Segment Length Comparisons

The AP lengths of the three major palatal segments (i.e., primary palate, anterior secondary palate, and posterior secondary palate) were estimated in millimeters from landmark coordinates without superimposition or scaling. To estimate the length of these segments along the anterior-posterior axis of the palate, the mean midline position between bilateral landmark pairs were calculated (Fig. 4). Proportional palatal segment lengths were calculated as the length of a single segment divided by the sum of all three segment lengths for a given specimen.

**Figure 4.**
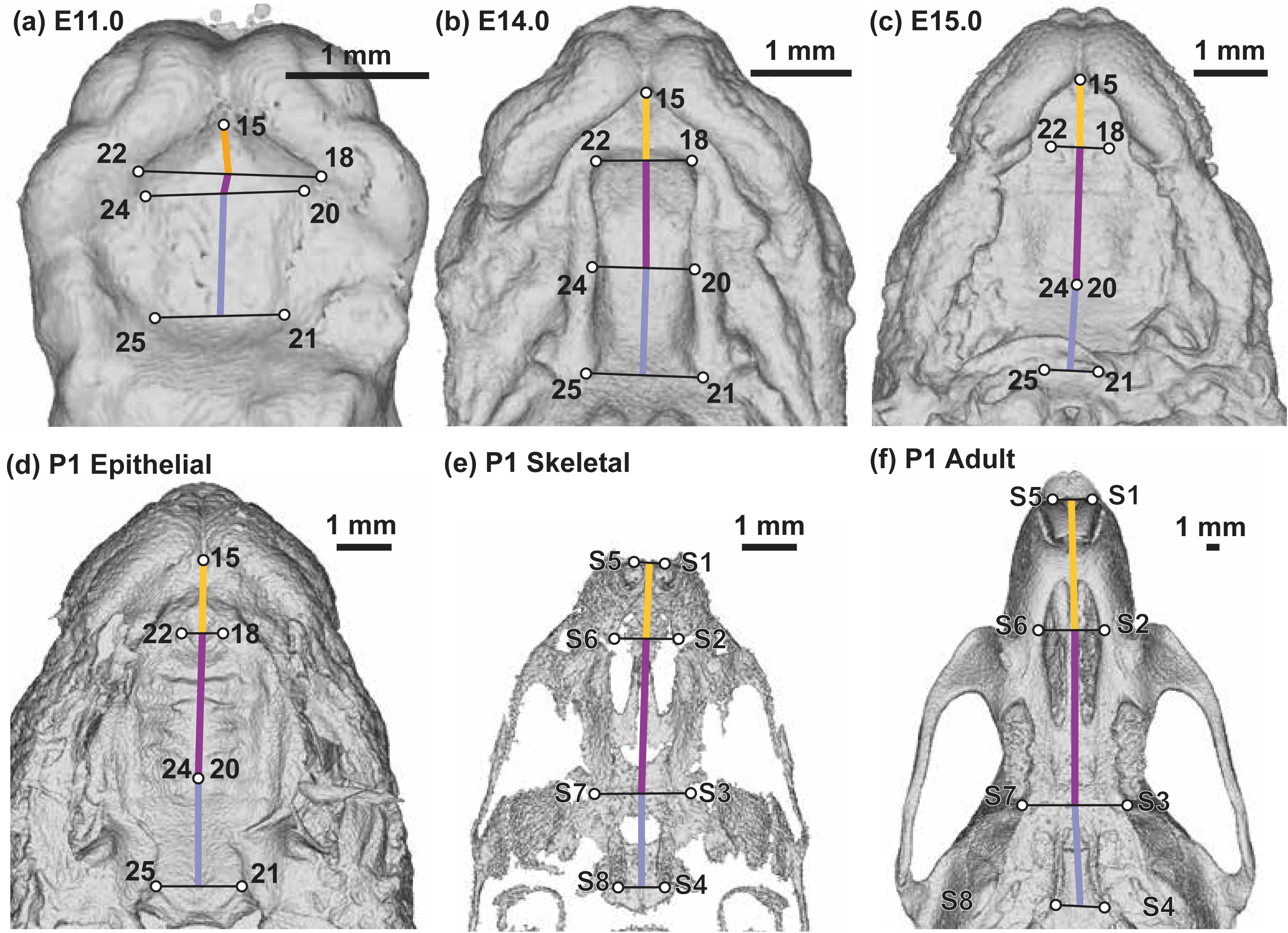
Palate segment length measurements – Identification of landmarks used to estimate midline lengths of the primary palate (yellow), anterior secondary palate (dark purple), and posterior secondary palate (light purple) on (a) E11.0, (b) E14.0, (c) E15.0 epithelial, (d) P1 epithelial, (e) P1 skeletal (after removing some neurocranial and facial bones to help with visualization), and (f) adult skeletal surfaces. Scale bars: 1mm. See also Supporting Tables 1 and 2.

Wilcoxon rank sum tests were used to identify significant differences in proportional palate segment lengths between age categories and between epithelial and skeletal P1 landmarks.

## Results

### Secondary Palate A-P Growth and the Position of Ruga 1 at a Facial Growth Center

Close relationships between different anatomical domains underlying midfacial outgrowth are found centered near ruga 1 (Fig. 5 & Supporting Fig. 2, red arrowheads). Our mouse embryonic series confirmed that ruga 1 formed within the MxP from a domain of *Shh* expression at the anterior extreme of the nascent secondary palate. As previously shown, other rugae then formed anterior to ruga 1 at the RGZ (Fig. 5c, asterisk) as the anterior secondary palate elongated, anteriorly displacing the primary palate and external nares. A separate domain of *Shh* expression posterior to ruga 1 gave rise to the geschmacksstreifen (gs) and sensory papilla arrayed across the most posterior palate. Our embryonic series also confirmed that ruga 1 is consistently found at the *Shox2/Tbx22* gene expression boundary between the anterior and posterior secondary palates during this critical embryonic period of palatal development (Fig. 5 & Supporting Fig. 2).

**Figure 5.**
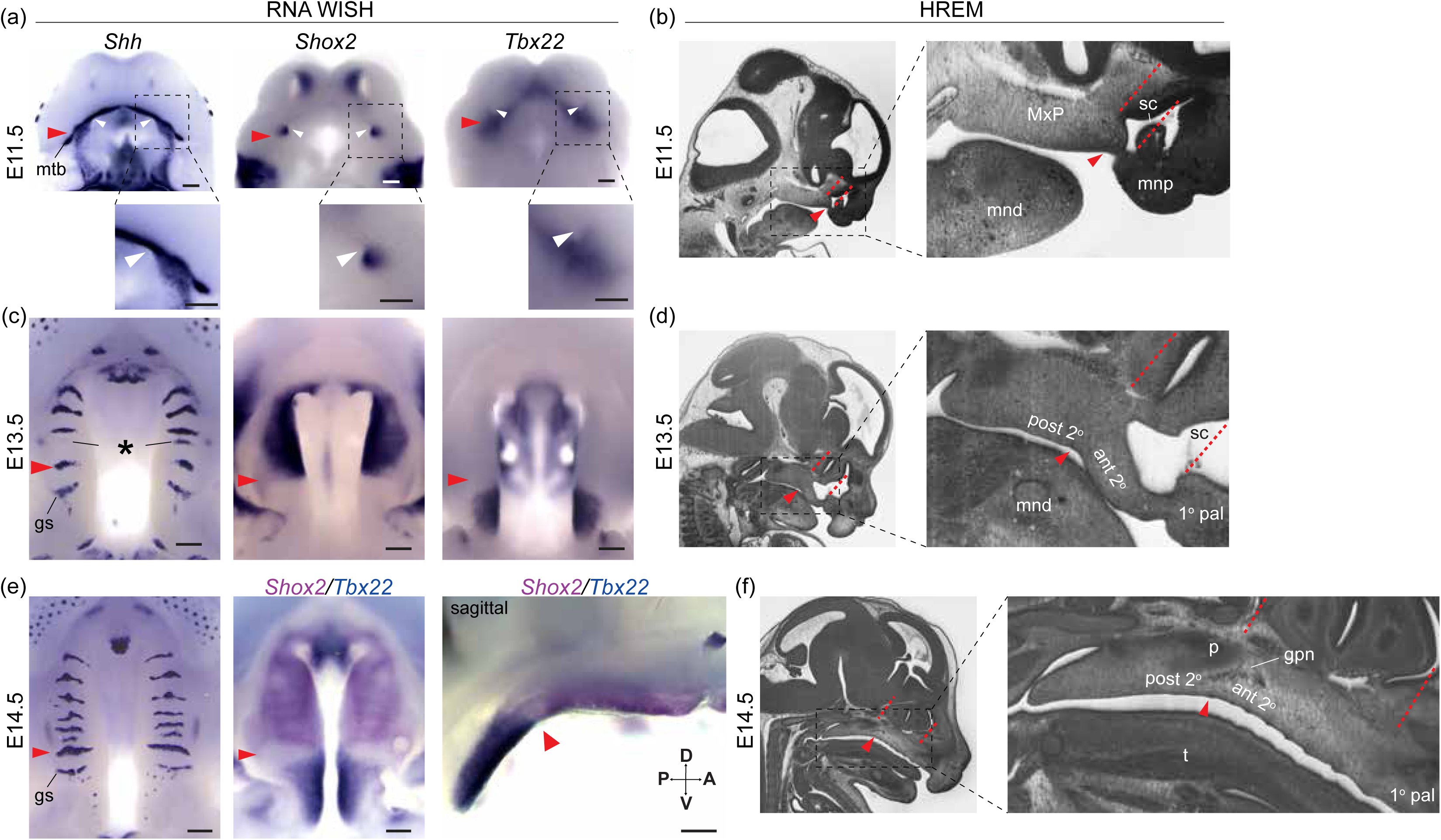
A-P molecular heterogeneity and anatomical relationships during secondary palate morphogenesis - (a, c, e) RNA WISH for *Shh*, *Shox2*, and *Tbx22* expression, and (b, d, f) sagittal sections of wildtype embryos imaged with high resolution episcopic microscopy (HREM) to provide histological resolution of the primary (1° pal), anterior secondary (ant 2°), and posterior secondary palate (post 2°) relative to surrounding facial structure during midfacial outgrowth at (a, b) E11.5 (c, d) E13.5, and (e, f) E14.5. Throughout midfacial outgrowth, ruga 1 (red arrowheads) marks the shared A-P expression boundary of *Shox2* and *Tbx22* in the anterior and posterior secondary palate, respectively. Double label WISH for *Shox2* (magenta) and *Tbx22* (blue) at E14.5 highlights mutually exclusive anterior and posterior expression domains organized relative to ruga 1 (oral view left, sagittal view right). White arrowheads mark the choanae, black asterisk marks the RGZ. Red dashed lines mark position of coronal planes passing through the primary-secondary palate junction and posterior wall of the nasal capsule to highlight coordinated elongation of the anterior secondary palate and overlying sinus cavity (sc). Regions in dashed boxes of (a) and in HREM images shown enlarged below or to the right, respectively. Additional abbreviations: mandible (mnd), molar tooth bud (mtb), maxillary process (MxP), medial nasal process (mnp), tongue (t), greater palatine nerve (gpn), geschmacksstreifen (gs), palatine (p), posterior domain of *Shh* expression (pd). Scale bars: 250um.

We identified a previously unappreciated association between ruga 1 and the posterior wall of the nasal capsule. At E11.5, ruga 1 formed adjacent to the primary choanae (Fig. 5a & Supporting Fig. 2a, white arrowheads), the posterior openings of the nasal passages, which initially occur at the boundary between the primary and secondary palatal segments (Tamarin, 1982). Elongation of the anterior secondary palate then coincided with the formation and expansion of the overlying nasal capsule. As the primary choanae and nasal capsule elongated between E11.5 and E15.5, ruga 1 and the posterior wall of the nasal capsule remained in approximately the same coronal plane (represented by position of red dashed lines in Fig. 5b, 5d, 5f and Supporting Fig. 2). Ruga 1 was also found coincident with a gross morphological inflection point of the palatal tissue where the anterior secondary palate and overlying nasal capsule exhibited an inferiorly accentuated angle relative to the cranial base, while the posterior secondary palate did not (Fig. 5d-f, Supporting Fig. 2).

The greater palatine neurovascular bundle was also consistently associated with ruga 1 across this embryonic period. The greater palatine nerve and artery were found entering the palatal shelves from a location immediately dorsal to ruga 1 at E12.5, and this spatial relationship was maintained until the greater palatine foramen formed by the articulation of maxilla and palatine bones around the palatine neurovascular bundle at the maxillary-palatine suture (Fig. 5f, Supporting Fig. 2). In combination, the consistent associations between ruga 1, the anterior-posterior secondary palate boundary, the posterior margin of the nasal capsule, and greater palatine neurovascular bundle suggested that these features are positioned at a proximal source of directional growth that the elongating anterior secondary palate and nasal capsule extend away from between E11.5 and E15.5.

A complementary 3D morphometric analysis of midfacial and palatal epithelial landmarks was used to quantify the integrated morphogenetic processes occurring during this critical developmental period. The first major principal axis of midfacial and palatal shape variation (PC1) for all limb bud staged E11-E15 specimens represented 73% of shape variance and was associated with general ontogenetic growth during this time window (Fig. 6). Mice with similar limb-based developmental age estimates fell near each other along the first principal component of shape variation. Additional principal axes of embryonic midfacial shape variation suggested shifts in the direction of ontogenetic changes over developmental time, with one inflection noted along PC2 (Fig. 6a), two along PC3 (Fig. 6b), and three along PC4 (Fig. 6c). Average developmental age category shape differences were illustrated as landmark specific ontogenetic shifts in palate shape (Fig. 7) and overall midfacial shape (Supporting Fig. 3).

**Figure 6.**
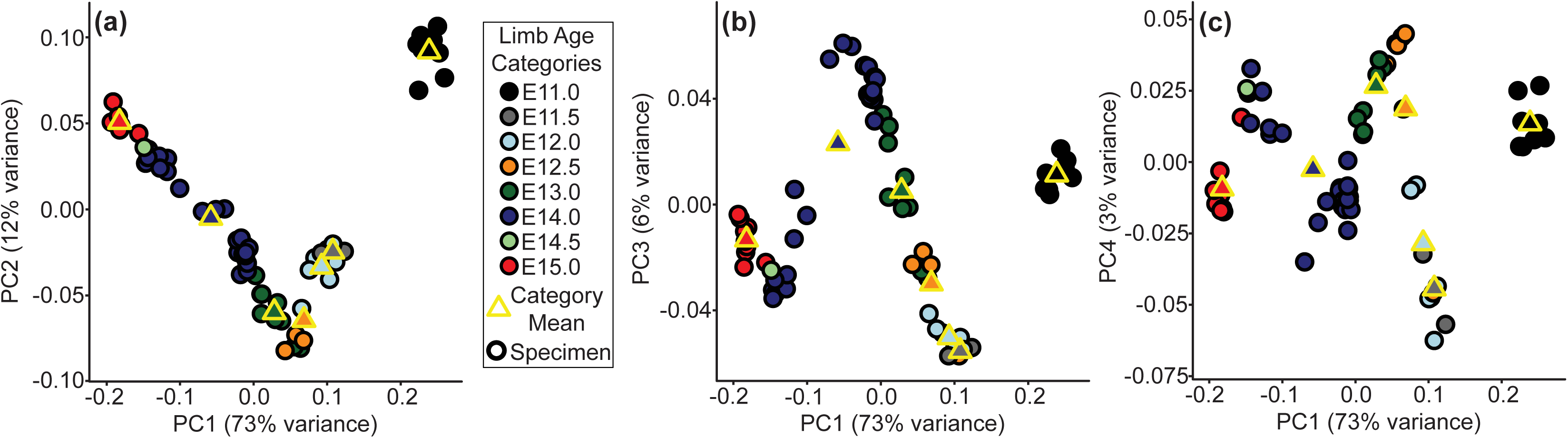
Major axes of embryonic midfacial and palatal shape variation - C57BL/6J embryo specimens plotted along the (a) first and second principal component axes of shape (i.e., PC1 and PC2), (b) along PC1 and PC3, and (c) along PC1 and PC4. Circle color indicates the limb based developmental age categories of each specimen, with the yellow bordered circles indicating the mean PC scores for each age category with more than one specimen. The proportions of midfacial and palatal shape variance associated with each principal component are provided.

**Figure 7.**
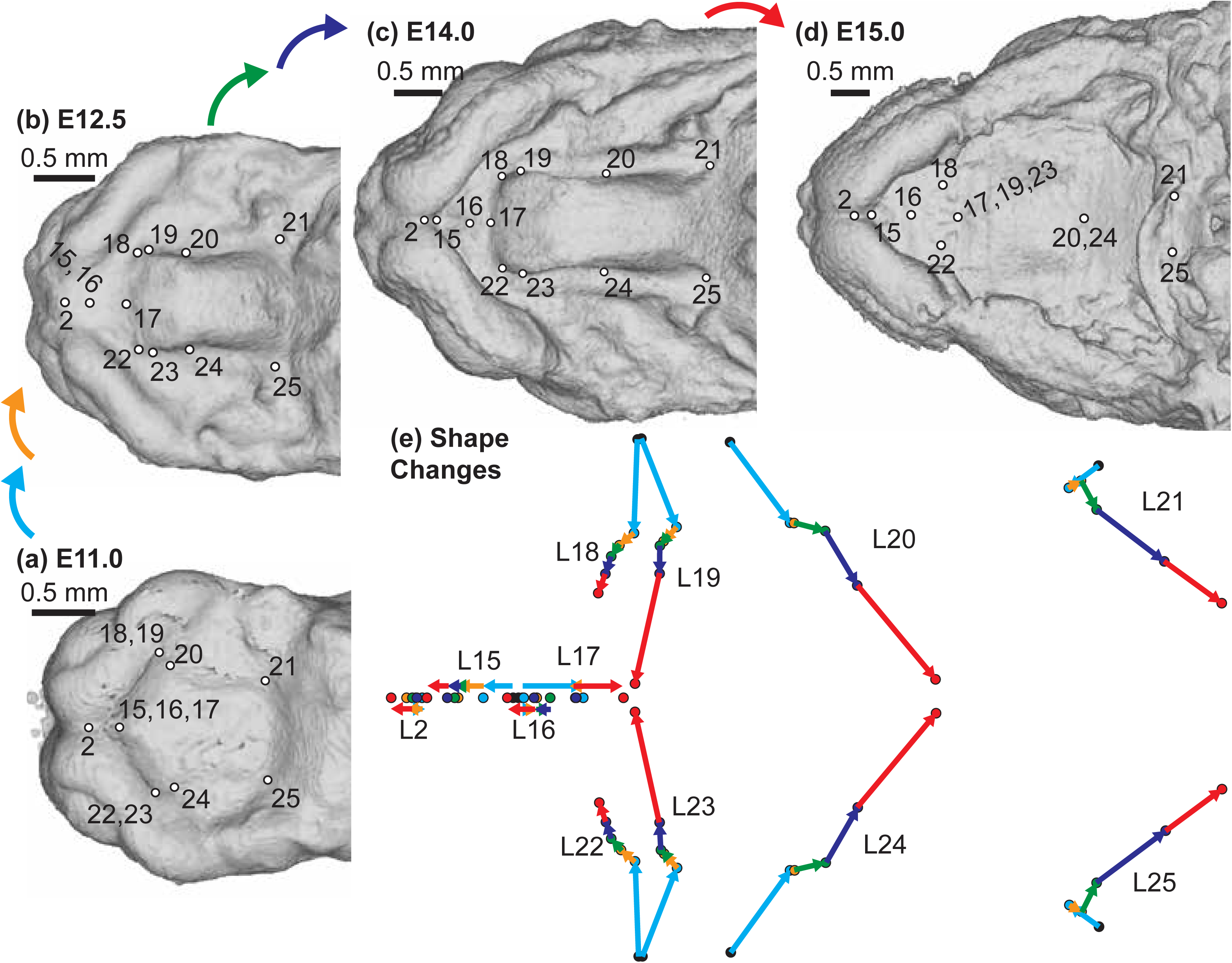
Palatal landmark growth trajectories – The positions of palatal landmarks are identified on representative C57BL/6J specimens at limb-derived developmental ages (a) E11, (b) E12.5, (c) E14, and (d) E15. (e) The average palatal landmark positions are plotted for each developmental age category that had more than one specimen. These landmark positions represent palatal shape after the removal of overall midfacial and palatal scale (as measured by centroid size) during Procrustes superimposition. The arrows indicate the trajectory of shape change for each landmark between ages. Black=E11; Light Blue = E12; Orange = E12.5; Green = E13; Dark Blue = E14; Red = E15. Scale bars: 0.5mm.

Between developmental stages E11 and E15, each of the three major palatal segments grew in length, with significant differences identified for almost all segment-specific consecutive age comparisons (Figs. 8a, 9a). However, the anterior secondary palate grew proportionally more than the primary palate and posterior secondary palate (Table 2, Figs. 8c, 9c). Soon after the start of embryonic A-P palate elongation, the primary palate represented ∼20% of the total palate length, the anterior secondary palate ∼20%, and the posterior secondary palate ∼60%. After E12.5, anterior secondary palate length increased proportionally faster than total palate length between each measured developmental day. By the time of midline palatal fusion, the primary palate represented ∼25% of palate length, the anterior secondary palate represented >40%, and the posterior secondary palate represented <35%. Therefore, between E11 and E15, the proportional contribution of the primary palate increased by a quarter while posterior secondary palate contribution decreased by nearly a third (Table 2, Fig. 9c). Strikingly, the proportional contribution of the anterior secondary palate to overall palate length doubled during this period.

**Figure 8.**
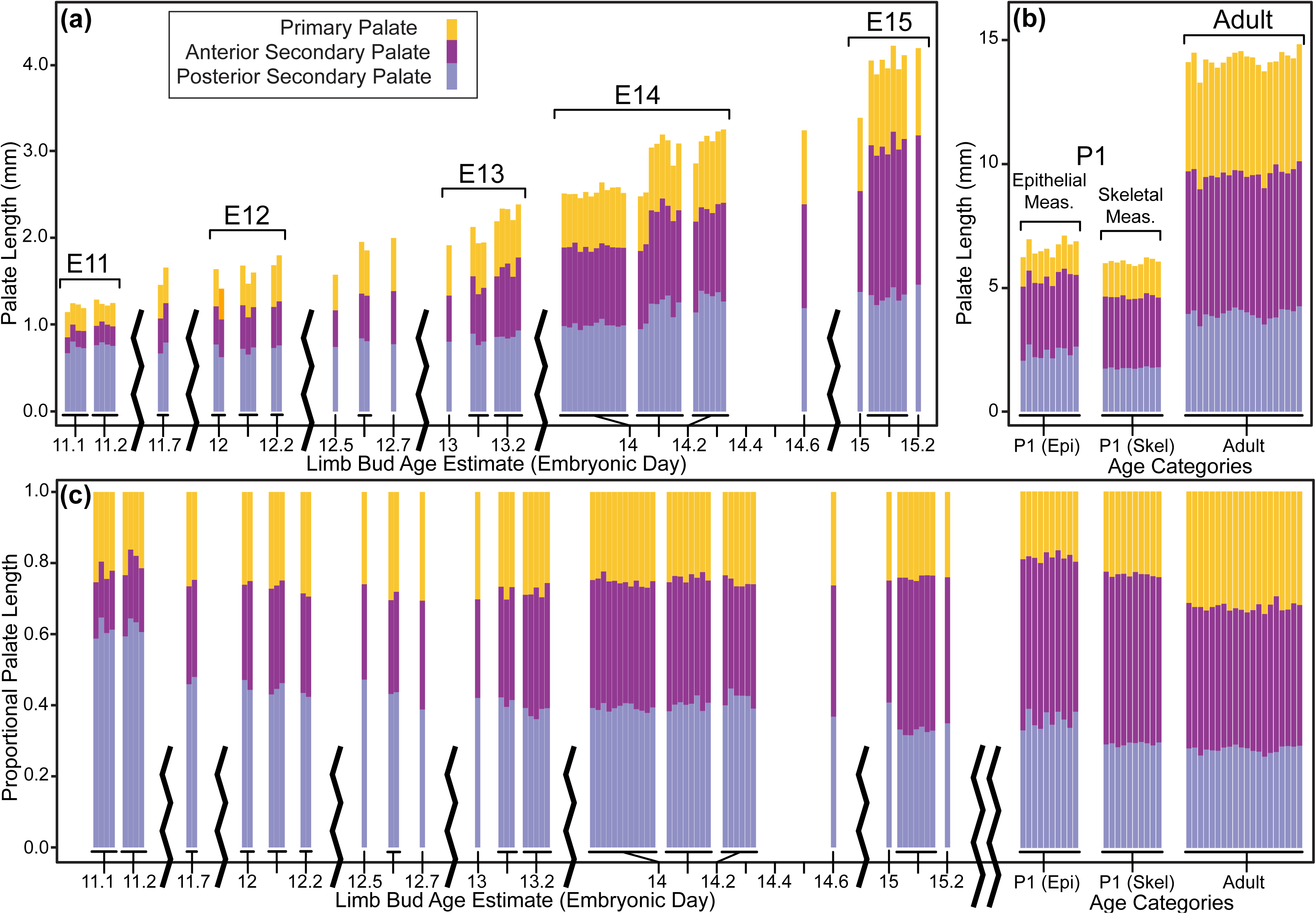
Palate segment length across ontogeny – The midline projected mean lengths (in mm) of the primary palate, anterior secondary palate, and posterior secondary palate for all (a) E11-E15 specimens and (b) postnatal specimens. (c) Palate segment length relative to overall midline palate length for the all specimens. Whole-day limb bud derived age categories and postnatal age categories used for pairwise statistical comparisons are indicated by brackets above the bars. Comparable segment lengths were collected from epithelial and skeletal surfaces for a single set of P1 specimens.

**Figure 9.**
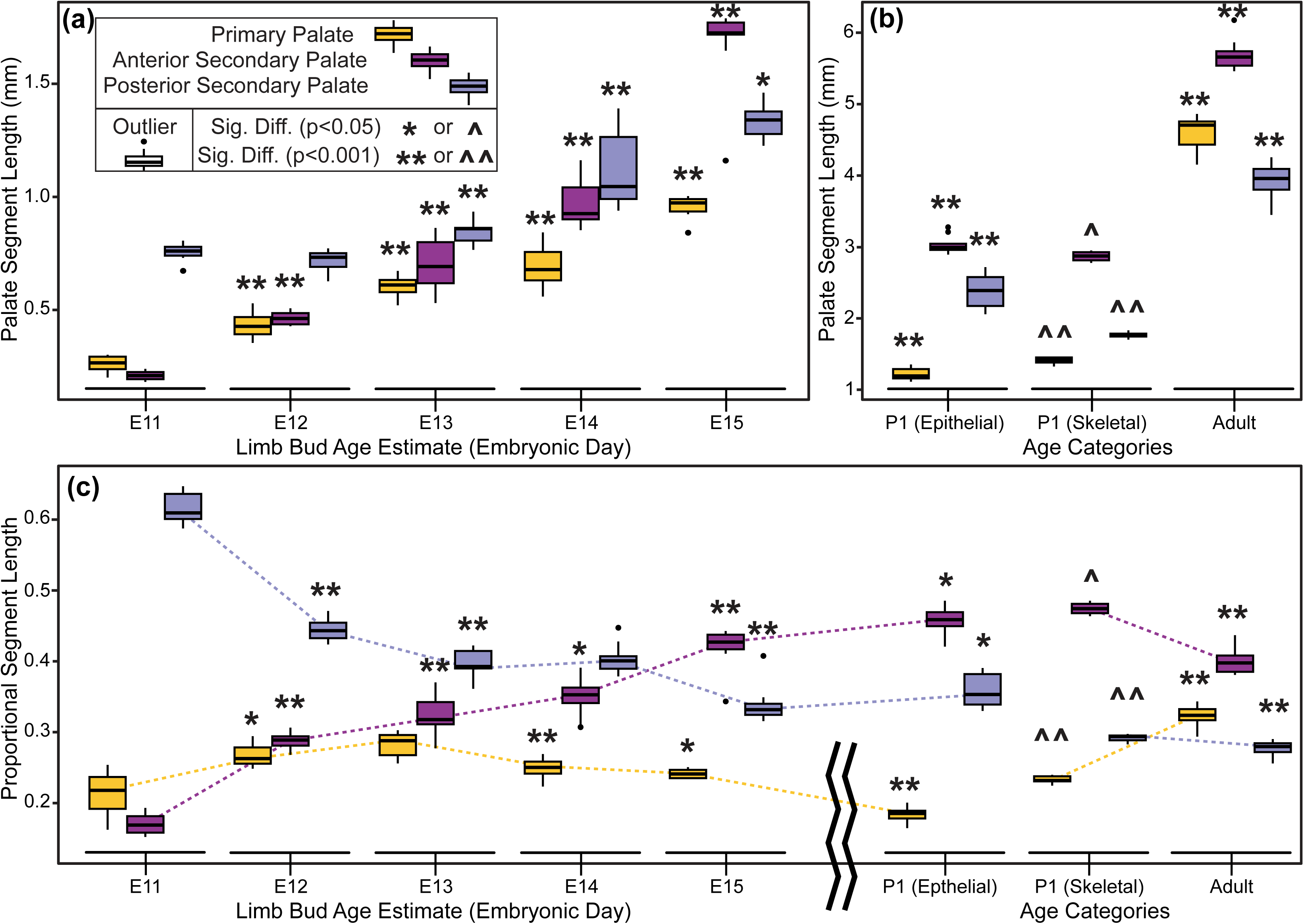
Palate segment length comparisons - Boxplots of palate segment specific lengths (in mm) for (a) whole-day limb dub derived embryonic age categories and (b) postnatal age categories. (c) Palate segment length relative to overall midline palate length for all specimens. Outliers are specimen values that are 1.5 times the interquartile range outside of the 25th or 75th quartile of that segment’s measures. Significant differences are indicated for each age that a given segment’s median value is different than the next younger age category (* when p-value is < 0.05; ** when p-value < 0.001). Significant differences for a given segment’s P1 skeletal measurement median when compared to the same segment’s P1 epithelial measurement (^ when p-value < 0.05; ^^ when p-value < 0.001). Dashed lines illustrate ontogenetic trends of palatal segment proportional contributions to overall palatal length.

**Table 2.** Median proportional palatal segment lengths for limb-based embryonic ages and postnatal age categories, presented as percentages of total palatal length with Q1 and Q3 in parentheses. Categories with enough samples for pairwise statistical comparisons are included. See **Fig. 9** for associated boxplots.

| Age Category and Tissue Measured | Sample Size | Primary Palate /Premaxilla | Ant. Sec. Palate / Maxilla | Post. Sec. Palate / Palatine & Pterygoid |
| --- | --- | --- | --- | --- |
| E11 epithelial | 8 | 21.8% (19.2, 23.7) | 16.9% (15.8, 18.2) | 60.9% (60.1, 63.6) |
| E12 epithelial | 7 | 26.3% (25.6, 27.9) | 28.9% (28.1, 29.4) | 44.3% (43.3, 45.5) |
| E13 epithelial | 9 | 28.8% (26.8, 29.6) | 31.8% (31.1, 34.3) | 39.3% (39.0, 41.5) |
| E14 epithelial | 26 | 25.0% (24.2, 25.9) | 35.3% (34.1, 36.3) | 40.1% (39.0, 40.7) |
| E15 epithelial | 9 | 24.1% (23.5, 24.7) | 42.7% (41.7, 43.8) | 33.2% (32.5, 34.0) |
| P1 epithelial | 10 | 18.5% (17.9, 18.9) | 45.9% (45.0, 46.9) | 35.3% (33.9, 38.2) |
| P1 skeletal | 10 | 23.2% (23.1, 23.8) | 47.4% (46.8, 48.1) | 29.4% (28.9, 29.6) |
| Adult skeletal | 20 | 32.4% (31.7, 33.3) | 39.8% (38.5, 40.8) | 28.0% (27.2, 28.4) |

### Segmental Differences in Skeletal Specification, Differentiation, and Growth During Secondary Palate Morphogenesis

We performed a WISH analysis of *Runx2*, *Sp7*, and *Phospho1* expression (Fig. 10) alongside double labeled WISH of *Shh* and *Sp7* between E12.5 and E15.5 (Fig. 11a) to characterize bone specification and differentiation in relation to A-P growth dynamics of secondary palate segments. The position of rugae, which provided an ordered set of temporally specific anatomical features, allowed for the identification of previously unappreciated associations between the palatal segments and developing palatal bones.

**Figure 10.**
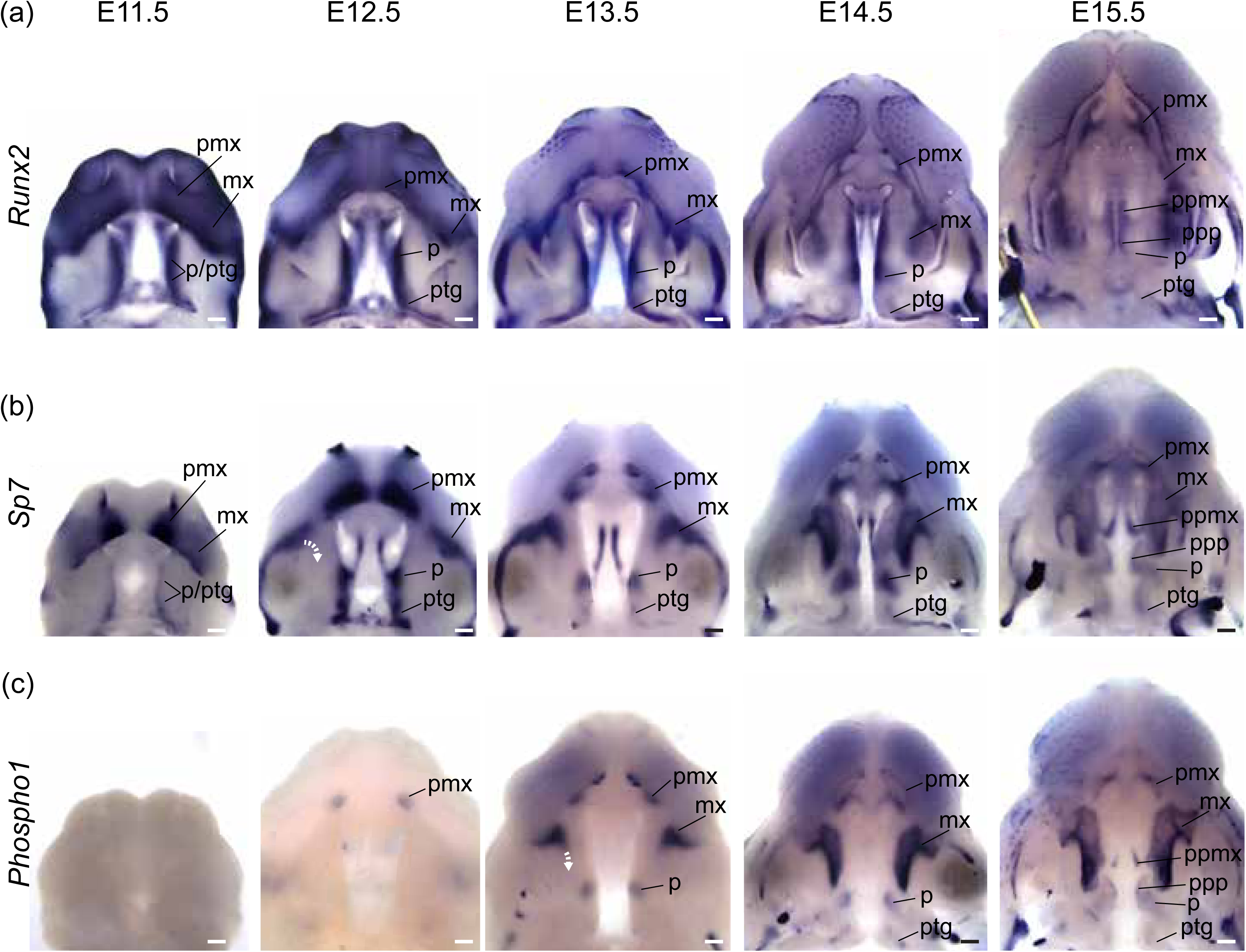
Segmental organization of skeletal specification and growth during A-P morphogenesis of the midface – WISH time course for (a) *Runx2*, (b) *Sp7*, and (c) *Phospho1* between E11.5 and E15.5. Expression of *Runx2* in osteoprogenitors highlights facial domains with osteogenic potential during midfacial morphogenesis. Expression of *Sp7* and *Phospho1* in committed and differentiating osteoblasts delineates the growth dynamics of individual skeletal anlagen during midfacial morphogenesis. Curved white dashed arrows in (b) and (c) indicate medial expansion of maxillary anlage. Abbreviations: maxilla (mx), palatine (p), premaxilla (pmx), pterygoid (ptg), palatal process of the maxilla (ppmx), palatal process of the palatine (ppp). Scale bars: 250um.

**Figure 11.**
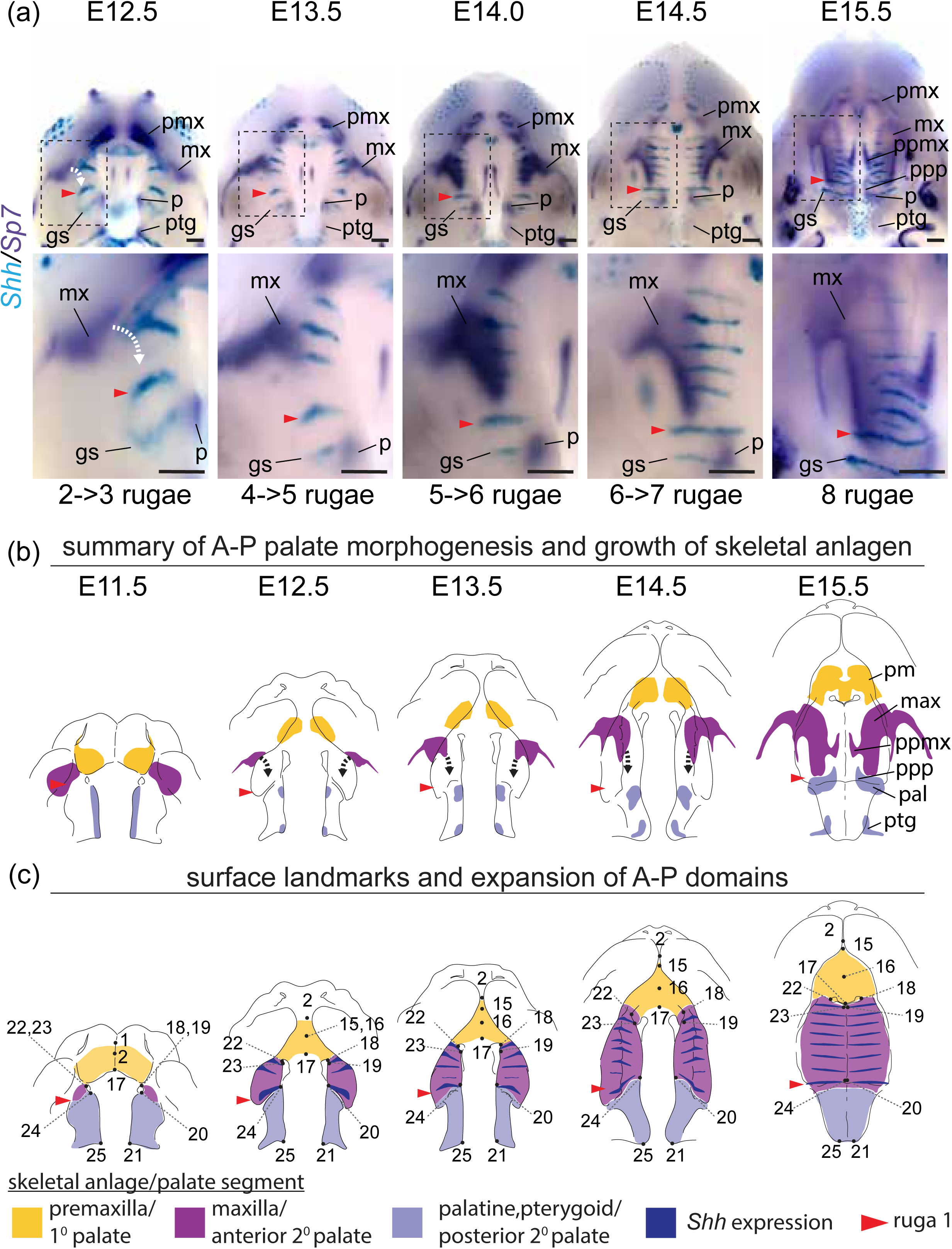
Double label RNA WISH time course for *Shh* (cyan) and *Sp7* (dark purple) between E12.5 and E15.5 - (a) Expression of *Sp7* in committed and differentiating osteoblasts delineates the growth dynamics of individual skeletal anlagen and *Shh* expression in rugae provides a temporally ordered set of A-P landmarks (regions in dashed boxes enlarged below). (b) Summary model of skeletal growth dynamics during midfacial outgrowth. Growth of the premaxilla (yellow) and palatine (pale blue) anlagen towards their characteristic shape occurs largely at the site of initial specification. Following initial specification external to the oral cavity, the maxilla (purple) grows into the anterior of the anterior secondary palate (curved white and black dashed arrows in a and b, respectively) towards the position of ruga 1 (red arrowhead) and palatine as expansion of the anterior secondary palate (double headed red arrow) separates the primary and posterior secondary palate. (c) Summary of the position of epithelial landmarks (see also Fig. 3) selected to capture segmental growth dynamics of the primary palate (yellow) and anterior secondary palate (purple) and posterior secondary palate (pale blue) during midfacial outgrowth. Abbreviations: geschmacksstreifen (gs), maxilla (mx), palatine (p), premaxilla (pmx), pterygoid (ptg), palatal process of the maxilla (ppmx), palatal process of the palatine (ppp). Scale bars: 250um

Broad *Runx2* expression in early facial prominences included osteogenic domains (e.g., premaxilla within the primary palate) and later became more restricted to regions adjacent to the molar tooth bud and medially along the adjoining palatal shelves (Fig. 10a). Osteoblast expression of both *Sp7* and *Phospho1* more precisely delineated the morphology of the emerging midfacial skeleton (Fig. 10b-c). Palatal processes of the maxilla and palatine bones formed within the medial region of the elevated palatal shelves, a domain that maintained high levels of *Runx2* expression during secondary palate growth (Fig. 10a), suggesting maintenance of a less differentiated osteoprogenitor population until palatal elevation and fusion was complete.

At E11.5, *Sp7* was strongly expressed in bilateral domains within the mnp of the forming primary palate. Between E12.5 and E15.5, presaging premaxilla morphology, *Sp7* expression formed a cup-like region surrounding the developing incisors (Fig. 10b). *Phospho1* was first observed in the forming premaxilla at E12.5 (Fig. 10c), reflecting a temporal lag between initial osteoblast commitment and later differentiation. The premaxilla acquired its characteristic adult morphology within the primary palate segment in which it was initially specified.

Within the secondary palate, this simple developmental sequence was repeated for the palatine and pterygoid bones within the posterior secondary palate, but development of the maxilla within the anterior secondary palate was unique. At E11.5, *Sp7* expression in the MxP was observed in two domains. The antero-lateral domain, which abutted the point of fusion with the mnp (i.e., the primary palate) and extended laterally away from the oral cavity and choanae, corresponded to the maxillary anlage. The posterior domain that spanned the length of the forming secondary palate, from the anterior limit abutting the choanae to the posterior-most palatal edge, gave rise to the palatine and pterygoid anlagen.

The antero-lateral domain of *Sp7* expression adjacent to the primary palate was initially external to the secondary palate at the site of the future zygomatic plate. However, this expression domain later expanded medially (E12.5-E13.5) and then posteriorly (E13.5-E15.5) within the growing anterior secondary palate (white curved dashed arrow in Fig. 10b). This expansion occurred after initial A-P growth of the anterior secondary palate and formation of 3-4 rugae anterior to ruga 1, suggesting that it is coupled with RGZ growth dynamics (Fig. 11a). The fact that *Phospho1* expression closely followed *Sp7* spatial dynamics further supported the idea that these *Sp7* expression patterns represent maxillary bone anlagen formation and expansion (white curved dashed arrow in Fig. 10c). The maxillary anlage continued growing posteriorly towards ruga 1 concomitant with rugae formation (white and black curved dashed arrows in Fig. 11a-b), within the elongating secondary palate.

The posterior domain of *Sp7* expression that initially spanned the length of the secondary palate at E11.5 was displaced posteriorly by expansion of the anterior secondary palate between E12.5 and E15.5. During this period, it separated into two subdomains. The anterior subdomain remained associated with the posterior wall of the nasal capsule and gave rise to the palatine bone, while the posterior subdomain formed both the medial and lateral pterygoid processes (Fig. 10b-c). Ruga 1 and the palatine anlage initially formed at the anterior extent of the secondary palate at E11.5 and maintained proximity to each other throughout palatal development even as both structures were displaced posteriorly (Fig. 11a).

The palatine and pterygoid anlagen grew to acquire their characteristic adult morphology within the posterior secondary palate segment where their precursor domains were initially specified, mirroring premaxillary anlagen formation within the primary palate. This differs from growth of the maxillary anlage that was initially specified external to the secondary palate at the site of the future zygomatic plate and only later extended into the anterior secondary palate, following a period of initial anterior secondary palate elongation, to give rise to the majority of the maxillary bone.

These results indicated that the posterior portion of the secondary palate and its associated anlage were present within the secondary palate at the onset of its morphogenesis at E11.5, while the anlage associated with the anterior secondary palate (i.e., the maxilla) did not contribute substantially to the secondary palate until later (Fig. 11b). They also indicated that the eventual meeting of the maxilla and palatine anlagen was an asymmetric process achieved predominantly by posterior growth of the maxilla towards the palatine (dashed black curved arrows in Fig. 11b). By E15.5, the maxilla and palatine anlagen met to form the maxillary-palatine suture subjacent to ruga 1 (see also Supporting Fig. 2).

Our results collectively revealed that the segmental relationship between three palatal regions and associated bone anlagen remained consistent throughout the period of palatal elongation, although the maxillary anlage was not present within the anterior secondary palate until partway through this period. Importantly, these results also highlighted the previously unappreciated correlation of maxillary growth to rugae formation as well as to the greater proportional expansion of the anterior secondary palate between E11.5 and E15.5 (summarized in Fig. 11c).

### Ontogeny of Postnatal Palatal Segment Length

We compared palatal segment lengths 1) between E15.5 and P1 based on epithelial surface landmarks and 2) between P1 and adult specimens based on skeletal landmarks of bony elements in 1:1 association with each palatal segment. Unsurprisingly, the length of each palate segment increased significantly between E15 and P1 based on epithelial measures, and between P1 and adult specimens based on skeletal measures (Figs. 8b, 9b). Proportional palate measures of all palate segments also changed significantly for the same age group comparisons (Table 2, Figs. 8c, 9c). Continuing the trend previously identified between E11.5 and E15.5, the anterior secondary palate grew proportionally more than the other segments between E15 and P1.

However, based on palatal bone measurements, the primary palate derived premaxilla elongated proportionally more than the other segments between P1 and adult specimens, (Table 2, Fig. 9c). The posterior secondary palate contribution remained stable (∼27-29%) across the postnatal period, but the primary palate contribution increased from ∼24% to ∼32% and the anterior secondary palate contribution decreased from ∼47% to ∼40% (Table 2). Although the A-P growth of the anterior secondary palate was the largest proportional contributor to midfacial growth between E11.5 and P1, outgrowth of the premaxilla plays a larger role during postnatal midfacial outgrowth.

## Discussion

Our results provided a multifaceted characterization of 1) normal murine midfacial growth dynamics within the secondary palate and upper jaw bones during the earliest stages of palatal growth and 2) subsequent changes in upper jaw bone proportions in newborn and adult mice. By tracking the position of the RGZ, sequential palatal rugae formation, and palatal bone precursor populations across the earliest phases of palatal A-P elongation, our results illustrated that the first-formed palatal ruga (i.e., ruga 1), which sits at an important border of regulatory gene expression, is coincident with important nasal, neurovascular and palatal structures throughout early midfacial development. For example, ruga 1 represents a consistent morphological boundary between the presumptive maxilla of the anterior secondary palate and the presumptive palatine and pterygoid bones of the posterior secondary palate. This association suggested that the process of rugae morphogenesis is coupled to maxillary osteogenesis during anterior secondary palate elongation, which contributes disproportionately to embryonic palate growth. It also verifies that ruga 1 approximates the future position of the maxillary-palatine suture from the time of its formation, rather than being associated with the boundary between the hard and soft palates.

### Normal Upper Jaw and Palatal Elongation

It is well known that A-P elongation of the palate is a necessary component of the integrated processes of midfacial outgrowth. However, within mammals, we previously lacked a detailed picture of the earliest spatiotemporal dynamics of intramembranous midfacial skeletal specification and differentiation within the context of the surrounding palatal segments and rugae formation. Our analysis indicated that the relationship between three longitudinal palate segments and associated upper jaw bones was already established during the earliest phases of palatal morphogenesis. These results illustrated the substantial contribution of embryonic anterior secondary palate elongation to overall palate length in mice and likely in other mammals. Anterior secondary palate elongation was also directly associated with rugae formation at the RGZ and elongation of the maxillary bone primordium.

Beyond verifying that ruga 1 forms at an important gene expression boundary between the anterior and posterior secondary palate, our results clarified that ruga 1 represented a stable morphological boundary positioned near the future maxillary-palatine suture. Ruga 1 formed at the earliest stages of anterior secondary palate elongation, then maintained a position at the anterior edge of the palatine bone anlage as it was posteriorly displaced by expansion of the anterior secondary palate. Anterior to ruga 1, the maxillary bone osteogenic population expanded from the site of initial specification (i.e., the zygomatic plate) into the secondary palate after ruga 1 was posteriorly displaced during anterior secondary palate elongation. Given that the secondary palate initially only contained osteogenic domains of the posterior secondary palate (palatine and pterygoid precursors), these results suggested that cellular cues in the proximity of ruga 1 may initially inhibit maxillary bone formation within the secondary palate. Additionally, expansion of the palatal portion of the maxillary anlage was uniquely associated with RGZ dynamics and anterior secondary palate elongation, suggesting that RGZ morphogenesis plays a major role in determining the proportional contribution of the maxilla to the upper jaw.

Differences in upper jaw suture anatomy may also be related to segmental differences in early bone growth. The premaxilla and maxilla were specified in adjacent tissue domains that maintained close apposition throughout palate morphogenesis, suggesting equal contributions to the formation of the premaxillary-maxillary suture. Conversely, the palatal portion of the maxilla grew posteriorly towards the palatine anlage at ruga 1, suggesting that maxillary growth (and likely RGZ dynamics) played a larger role than palatine growth in determining maxillary-palatine suture position. Different growth dynamics at these sutures may lead to a vertical premaxillary-maxillary suture versus an oblique maxillary-palatine suture with substantial A-P overlap of the maxilla and palatine bones.

Because suture formation provides a critical niche for skeletal progenitors (Zhao et al., 2015), the way palatal segments contribute to palatal sutures may contribute to variation in postnatal dynamics of midfacial growth and remodeling (Enlow and Bang, 1965; Kurihara et al., 1980; Sarnat, 1997; Martinez-Maza et al., 2013; Vora et al., 2015; Maga, 2016). The widespread presence of Gli1+ mesenchymal stem cells and nascent bone within the adult maxillary-palatine suture and the maxillary bone, but not in the palatine bone (Luo et al., 2019) provides further support for this hypothesis. Given the established role of calvarial sutures in directing postnatal craniofacial bone growth, further comparisons of osteoprogenitor dynamics during premaxillary-maxillary and maxillary-palatine suture formation are warranted.

### Palatal Growth as a Basis for Morphological Variation

Given its central contribution to total palatal elongation, variation in RGZ-regulated A-P growth of the anterior secondary palate may play a role in determining the proportional contribution of the maxilla to the upper jaw and contribute substantially to the range of prognathism observed amongst mammals (Young et al., 2014). Support for this idea comes from the fact that more prognathic species tend to have more rugae: flat-faced humans typically have 3-6 palatal rugae (Hauser et al., 1989; Jayasankar et al., 2016), mice have 8-9 rugae (Peterkova et al., 1987), and prognathic pigs have 20-25 rugae (Tonge and McCance, 1965). Variation in interspecies rugae number and palatal segment proportions may stem from ontogenetic shifts in the relationship of various midfacial structures or shifts in the relative timing of normal developmental events occurring within different parts of the face.

This hypothesis could be tested within an evo-devo context by characterizing variation in palatal *Shh* expression between mammals and avians that have drastic differences in maxillary morphology. These differences include MxP-driven palatal growth in mammals and FNP-driven palatal growth in birds (Young et al., 2014) as well as an obligate avian cleft palate.

Alternatively, the ontogenetic basis for variation in palatal segment contributions could be dissected genetically via analysis of *Shh* expression and rugae morphogenesis in mice with altered A-P skeletal patterning, such as the *Pbx* CNCC mouse mutants (Welsh et al., 2018). Although changes within the RGZ growth dynamics of anterior secondary palate growth may be important for major species-specific differences in prognathism, postnatal growth processes also likely play an important role in determining adult upper jaw bone proportions.

### Concluding Statement

Our multifaceted illustration of normal midfacial growth dynamics confirmed a one-to-one relationship between palatal segments and upper jaw bones during the earliest stages of palatal growth, suggesting that the first formed ruga represents a consistent morphological boundary between anterior and posterior secondary palate bone precursors, thus approximating the position of the future maxillary-palatine suture. In addition to driving rugae formation, our results suggested that interactions at the RGZ coordinate elongation of the maxillary bone primordium within the anterior secondary palate, which more than doubles in length prior to palatal shelf fusion. Overall, our integrated analysis of spatial gene expression dynamics and palatal ontogenesis during midfacial elongation provides a critical new perspective within which to consider palatal segment contributions to both normal developmental processes and the etiological basis of genetic perturbations.

## Acknowledgements

We are indebted to Benedikt Hallgrímsson for the use of his laboratory for uCT and microscopy image collection. This work was supported by start-up funds to Christopher J. Percival from Stony Brook University; NIH/NIDCR R01 grant DE028753 and UCSF Chancellor Funds from Recruitment Package to Licia Selleri who supported the personnel and all the research completed in her lab for this study; and NIH/NIDCR F32 Fellowship DE026950 to Ian Welsh. The authors of this manuscript have no conflicts of interest to declare.

## Data Availability Statement

µCT images, limb bud microscopy photographs, landmark coordinate data, and associated morphometric analysis R scripts will be available publicly as a dataset within Dryad (https://doi.org/10.5061/dryad.ghx3ffbvb; Welsh et al., 2024) when this manuscript is accepted for publication.

## Author Contributions

ICW contributed to study design, acquisition and interpretation of histological images, and drafting of the manuscript. MEF contributed to acquisition, analysis, and interpretation of morphometric data, and critical revision of the manuscript. DL contributed to the design of morphometric methods, acquisition of morphometric data, measurement and analysis of limb bud images, and critical revision of the manuscript. IM contributed to acquisition of µCT images, acquisition, measurement and analysis of limb bud images, and critical revision of the manuscript. KH contributed to mouse breeding, specimen collection, histological data collection, and critical revision of the manuscript. CJP contributed to study design, acquisition of µCT images, acquisition, analysis, and interpretation of morphometric data, and drafting of the manuscript.

---

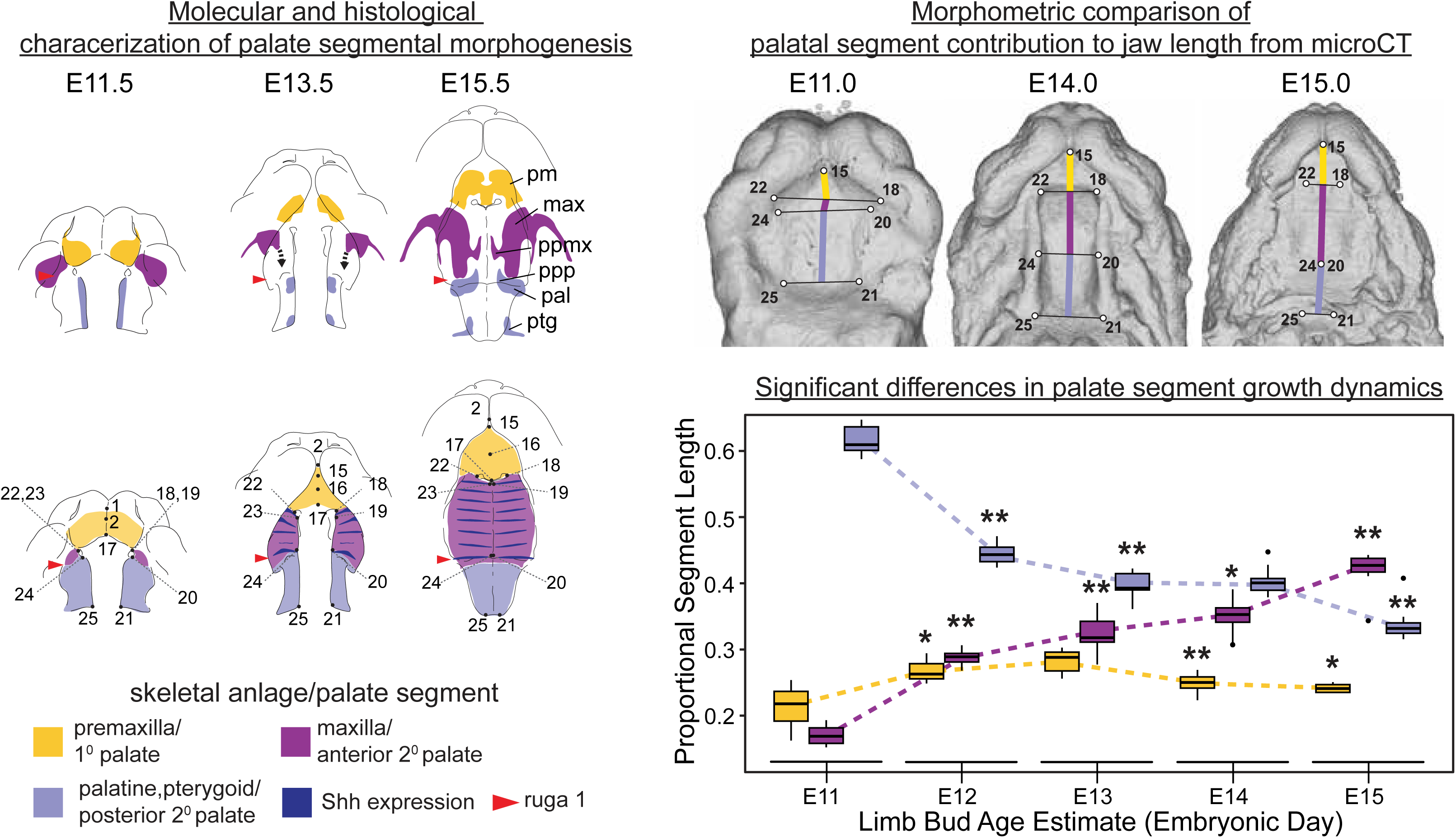

## Supporting Information

Supporting Tables and Figures are found as a Supporting .pdf file in association with this manuscript on the publisher’s website.

