## Supporting Tables and Figures for "Palatal segment contributions to midfacial anterior-posterior growth"

Supporting Table 1 – Epithelial landmark definitions, collected on the superficial tissue surface of the face and palate of developmental age E11 – E15 specimens, with a subset collected on P1 specimens. See also Fig. 3.

| Landmark Number | Epithelial Landmark Definition |
|---|---|
| **1** | Most rostral midline point at the meeting of the medial nasal processes, becoming the most rostral midline point on the rhinarium. |
| **2** | The most ventral midline point along the developing lip on the primary palate, in the region that develops into the lip vermillion. |
| **3(9)** | Dorso-caudal corner of the whisker field, taken on the skin right next to the plateau of the whisker field, rather than on the field itself; at E11.5, taken where the lateral nasal process and forebrain region meet. |
| **4(10)** | Rostral apex of the forming medial canthus of the eye; not collected at P1 because of eyelids. |
| **5(11)** | Caudal apex of the forming lateral canthus of the eye, not collected at P1 because of eyelids |
| **6(12)** | The caudal most point of the intersection between the lateral nasal process and the maxillary process; after E13, taken at the edge of the margin between second and third whisker rows (counting from the top). |
| **7(13)** | Point at the middle of the medial side of the nasal aperture as a lateral extent of the medial nasal process, taken on the medial edge of the nasal aperture at the point of concave inflection between upper and lower portions of the developing nasal aperture. |
| **8(14)** | The dorso-lateral most point of the nasal aperture |
| **15** | Most rostral midline point of the primary palate ruga; at E11.5, it may be the same position as LM 16 and 17. |
| **16** | Most caudal midline point of the primary palate ruga; at E11.5, it may be the same position as LM 15 and 17. |
| **17** | Most caudal midline point of the primary palate; at E11.5, it may be the same position as LM 15 and 16. |
| **18(22)** | Rostro-buccal extent of secondary palate, placed at a lateral edge of the secondary palate where it meets the primary palate; at E11.5, it may be at the same position as LM 19(23). |
| **19(23)** | Rostral-lingual corner of the palatal shelf, placed on the rostro-medial corner of the growing secondary palatal shelf; at E11.5, it may be at the same position as LM 18(22). |
| **20(24)** | Medial-most palatal shelf point on the border between anterior and posterior secondary palate, taken at the dorsal ventral inflection of the palatal shelf. |
| **21(25)** | Caudal-lingual corner of the palatal shelf; after E13 the point is taken in proximity to the posterior end of the pterygoid process (i.e., not on the posterior end of the pterygoid process, but at the caudal-lingual corner of the palatal shelf that is close to the posterior end of the pterygoid process). |

Supporting Table 2 – Skeletal palate landmark definitions collected on upper jaw bones of P1 and adult specimens. See also Fig. 4.

| **Epithelial Homolog** | **P1 & Adult Number** | **Definition** |
|---|---|---|
| 15 | S1(S5) | Most rostral point of the superior incisor alveolus, taken at the midline of the incisor. In P1, taken at approximate location of the incisor midline due to incomplete dental development. |
| 18(22) | S2(S6) | Ventral point on the maxilla-premaxillary suture, taken on the premaxilla. |
| 20(24) | S3(S7) | Most caudal point on the molar alveolus, taken on approximately the midline axis of the molar row. In P1, taken at the most caudal point of the developing maxillary alveolus. |
| 21(25) | S4(S8) | Most ventrocaudal point of the pterygoid process. |

### Supporting Figures

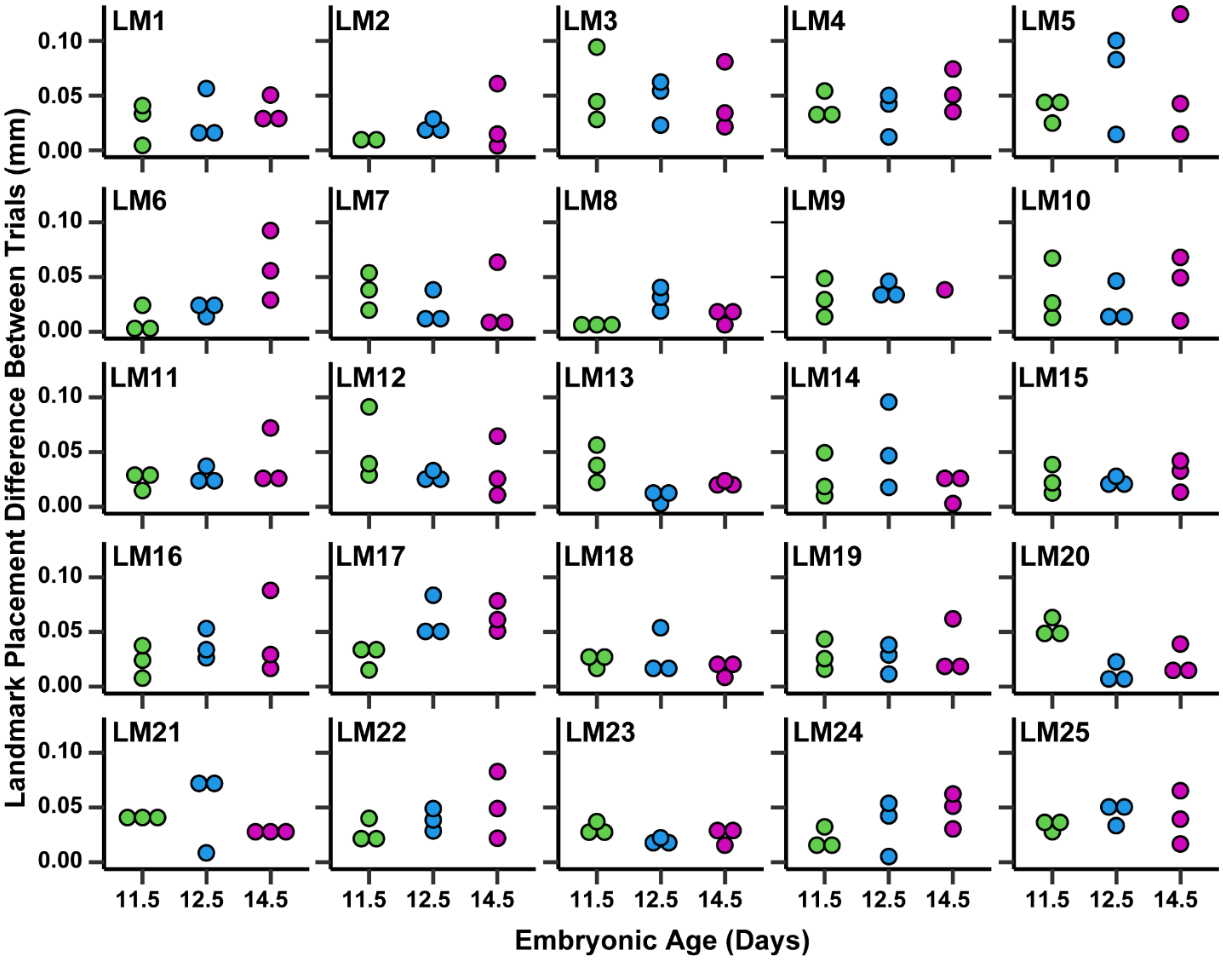


Supporting Figure 1 – Intra-observer landmark placement error - Difference in landmark placement between trial 1 and trial 2 collected for the same nine specimens (3 E11.5, 3 E12.5, and 3 E14.5) by a single observer. Distance between landmark positions of specific landmarks in the two trials is measured in millimeters.

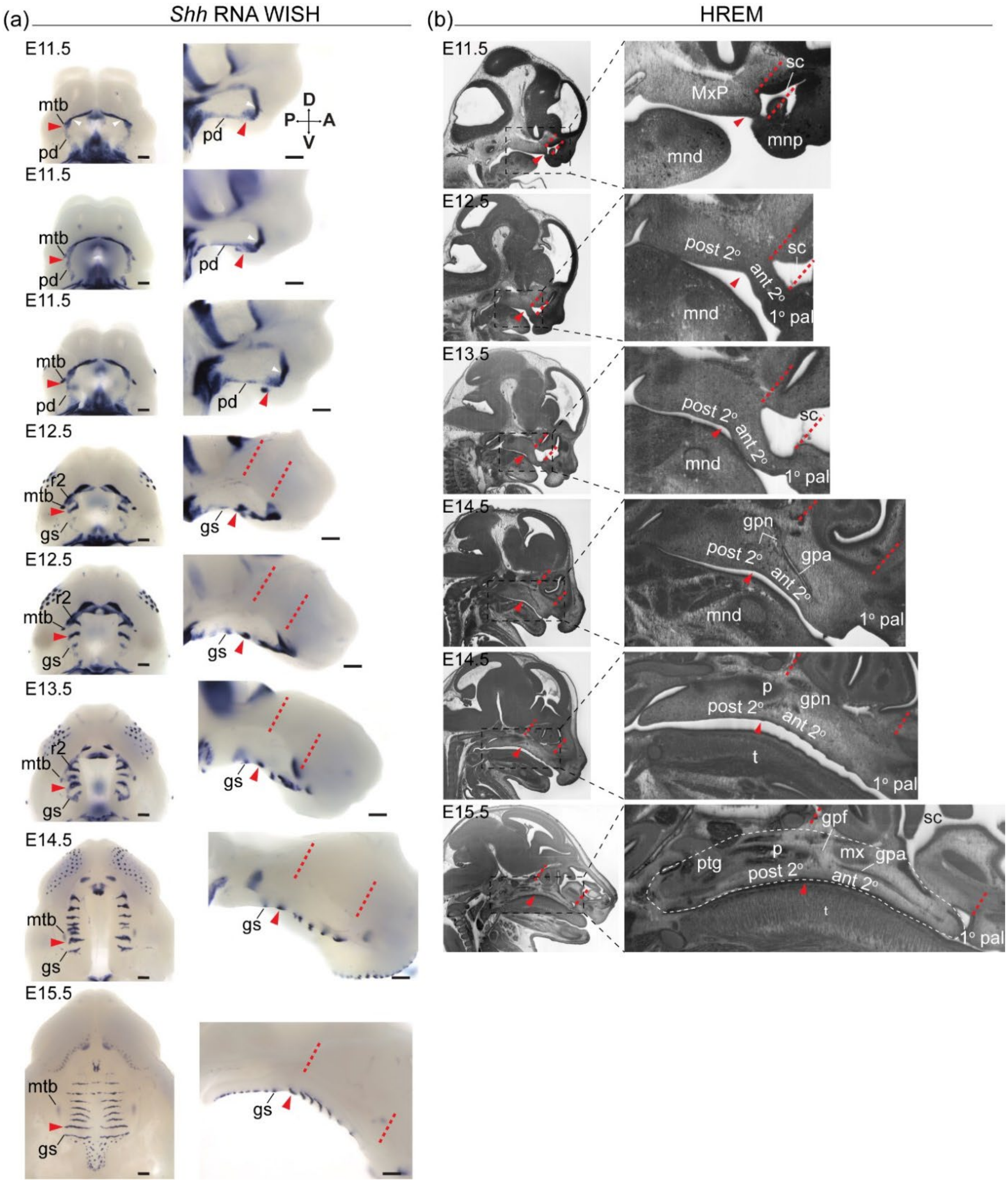


Supporting Figure 2 – Segmental growth dynamics of the midface of E11.5 through E15.5 wildtype embryos relative to A-P landmarks visualized via (a) RNA WISH for *Shh* expression or (b) high resolution episcopic microscopy (HREM). (a) RNA WISH for *Shh* in the oral cavity viewed orally (left column) or sagittally (right column) highlights dynamics of ruga 1 formation from an anterior domain adjacent to the choanae the relative to position of the forming molar tooth bud laterally, ruga 2 anteriorly, and a posterior domain that will form the geschmacksstreifen. (b) Sagittal sections imaged with provide histological resolution of anatomical landmarks A-P orientation of during midfacial elongation between E11.5 and E15.5. Two E14.5 samples show pre- and post-elevation of the secondary palate. Red arrowhead marks the position of ruga 1, white arrowheads mark the choanae, red dashed lines mark position of coronal planes passing through the primary-secondary palate junction and posterior wall of the nasal capsule to highlight coordinated elongation of the anterior secondary palate and overlying sinus cavity. White dashed outline demarcates the secondary palate at E15.5. HREM data available at: https://www.ebi.ac.uk/biostudies/bioimages/studies/S-BIAD490?query=DMDD.

Abbreviations: anterior secondary palate (ant 2$^{o}$), geschmacksstreifen (gs), greater palatine artery (gpa), greater palatine foramen (gpf), greater palatine nerve (gpn), mandible (mnd), maxilla (mx), maxillary process (MxP), medial nasal process (mnp), molar tooth bud (mtb), posterior domain of *Shh* expression (pd), posterior secondary palate (post 2$^{o}$), primary palate (1$^{o}$ pal), sinus cavity (sc), tongue (t). Scale bars: 250um.

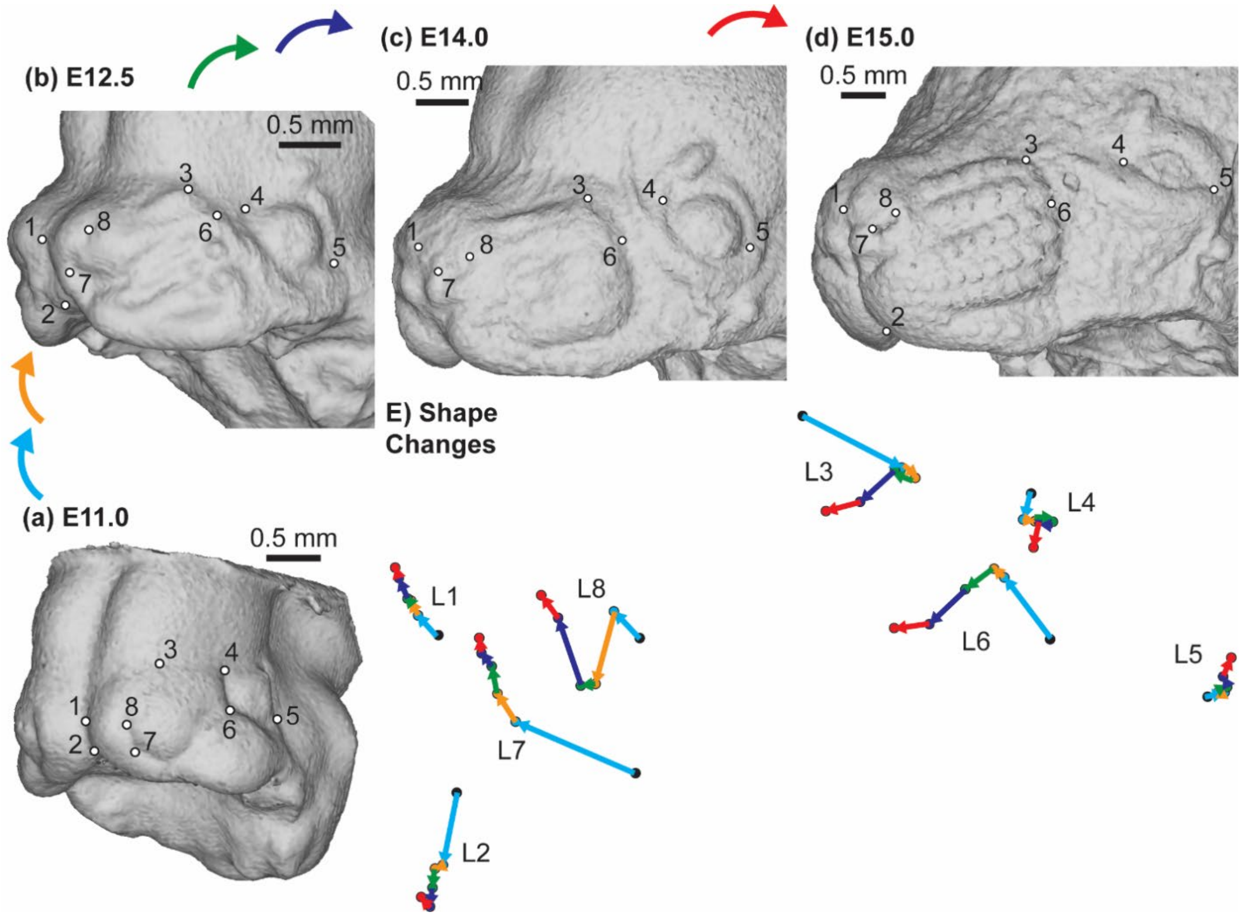


Supporting Figure 3 – C57 midfacial landmark growth trajectories – The positions of midfacial landmarks are identified on representative C57 specimens at limb-derived developmental ages (a) E11, (b) E12.5, (c) E14, and (d) E15. E) The average midfacial landmark positions are plotted for each developmental age category that has more than one C57 sample. These landmark positions represent midfacial shape after the removal of head scale during Procrustes superimposition. The arrows indicate the trajectory of shape change for each landmark between ages. Black=E11; Light Blue = E12; Orange = E12.5; Green = E13; Dark Blue = E14; Red = E15. Scale bars representing 0.5mm are provided for reference.

The most noticeable change in midfacial shape occurred between E11 and E12, as the facial prominences finished fusing and the nasal region began to elongate. During this period, the posterior edge of the whisker field consolidated and was found close to the anterior canthus of the eye. However, the direction of shape vectors changed after E12.5, as the relative distance between the whisker region and eye increased. In addition, changes in shape vectors were noted for two landmarks of the nasal aperture (L7 and L8) as the shape of the nostril remained relatively short and bent with the growth of nearby portions of the rhinarium.

The double shift of specimen PC3 scores across developmental time (Fig. 6b) were partially based on these nasal shape changes.
